# Polymer Model Unveils Quantitative Association of Chromatin Conformation and Gene Regulation

**DOI:** 10.1101/2023.12.04.569923

**Authors:** Swayamshree Senapati, Inayat Ullah Irshad, Ajeet K. Sharma, Hemant Kumar

## Abstract

The spatial organization of chromatin in the nucleus is of essence for regulating gene transcription. However, the mechanisms governing the intricate interplay of chromatin structure and gene transcription remain poorly understood. Hi-C experiments have unveiled a multiscale chromatin organization, significantly enriching our understanding of the structural control of gene expression. We introduce a computational framework to link chromatin structural modifications to gene regulation. This framework generates an ensemble of three-dimensional conformations of a given genomic locus using a bead-spring polymer model where the cHiC contact map is used as an input. We then correlate such chromatin conformations to the transcription level of the encoded genes using a Markov chain-based model, where binding/unbinding rates extracted from the molecular dynamics trajectories are used. By considering a specific example, we have demonstrated that the deletion of the CTCF binding domain between two consecutive TADs leads to a significant change in the enhancer-promoter interaction and gene transcription of the encoded genes, namely sox9 and kcnj2, responsible for limb development. Such a change in gene expression level is quantitatively consistent with experiments. Further insight from the polymer-based 3D conformation reveals that the higher gene expression level of the kcnj2 gene is caused by the specific enhancers of kcnj2, present in the sox9 TAD, which become accessible after boundary deletion. By quantifying the impact of these enhancers, our model can also be used to identify the functional enhancers. Together, the present computational framework not only advances our understanding of the relationship between the spatial architecture of the chromosome and the function of the cell but also provides invaluable insights into potential therapeutic interventions targeting aberrant gene regulation in pathological contexts.

## 1. Introduction

The regulation of gene expression is controlled by various elements, including cis-regulatory elements (CREs) such as enhancers and promoters, as well as trans-regulatory elements (TREs) like transcription factors (TFs) and coactivators [1]–[3]. Although genes and their regulatory elements may be scattered over long genomic distances, they physically interact in three-dimensional space to regulate gene expression. The specific contacts are achieved through genome organization, which brings together the regulatory elements associated with a target gene [4], [5]. Any deviation from the optimal structure can alter the way regulatory elements interact, potentially leading to the misexpression of genes. Enhancers may disrupt their contact with natural promoters and exhibit aberrant interactions with alternative promoters, resulting in the dysregulation of gene expression. Furthermore, the involvement of structural proteins such as CTCF and cohesin is crucial in facilitating the connection between enhancers and promoters, and thus the disruption of these proteins can lead to gene aberrations [6]–[9]. Nevertheless, quantitative understanding of the variability in gene expression levels attributed to genome organization remains largely unknown due to experimental limitations such as the limited resolution of HiC, and the lack of multiple probes to simultaneously detect multiple genomic loci, and the phototoxicity of fluoroprobes in microscopy and in-situ fluorescence hybridization techniques [10], [11]. Advanced modeling approaches can be beneficial to elucidate the role of specific arrangements of chromatin in gene regulation.

The physics-based chromatin models have served as important tools for understanding the intricacies of chromatin organization and contact between genetic elements within the nucleus. Drawing inspiration from principles of polymer physics and statistical mechanics, these models integrate experimental data on HiC and diverse omics data to simulate the dynamic spatial arrangement of chromatin segments within the nucleus [12]–[15]. To elucidate the relationship between gene expression and genomic organization, we developed a computational framework that utilizes the cHiC contact map and genomic positions of enhancers and promoters as input to compute kinetic parameters of the chromatin dynamics and make quantitative predictions of corresponding gene transcription levels. In principle, such models can also be used to explore how epigenetic marks, histone modifications, and chromatin remodelers collectively orchestrate the accessibility of transcriptional machinery to specific genomic loci, thereby modulating gene expression patterns.

One of the experimental approaches to elucidating the relationship between genome structure and function is through the rearrangement of topological associating domain (TAD) boundaries. Deletions, inversions, or duplications of TAD boundaries are often created to disrupt the genome organization and then quantify the respective gene misexpression [16], [17]. In this study, we take a specific example of TAD boundary deletion to modify the genomic structure, and apply our computational framework to predict the gene expression level corresponding to this change. Within our computational framework, first we construct a polymer model capable of providing us with a three-dimensional configuration of the chromatin with a cHiC contant map as input. Utilizing this polymer model, we were able to discern alterations in enhancer-promoter (E-P) interactions following the deletion of TAD boundaries. Then, by combining the simulated trajectory with a kinetic model of gene expression, we accurately quantified the variations in gene expression resulting from modifications in chromatin structure. Notably, our analysis established the specificity of enhancers in regulating gene expression and revealed the importance of chromatin organization in modulating specific E-P interactions in governing gene expression. We also demonstrate that the specific E-P interaction leads to dissimilar changes in the expression levels of two genes situated within adjacent TADs. Our framework emerges as a means to establish a quantitative link between chromatin structure and the expression of encoded genes. By capturing the non-linear relationships inherent in the complex regulatory network, our modeling framework will enhance our fundamental understanding of the impact of genome architecture on gene regulation.

## 2. Methods

We have developed a multistep computational framework to predict the relative gene expression level corresponding to the organization of chromatin characterized by a given cHiC contact map. The framework utilized a combination of cHiC contact maps and the genomic positions of enhancers and promoters as inputs. The first step of the approach determines the three-dimensional chromatin conformation of the genomic region of interest through a polymer model of chromatin. We employed molecular dynamics simulations to generate ensembles of three-dimensional chromatin conformations of the 6Mb sox9-kcnj2 locus of chr11 in wildtype (WT) and deleted CTCF (DELC) cells of the mouse limb bud cells (see SI, sections 1-3). We used a bead-spring polymer model, which describes the polymer as beads connected by springs [18]. Our polymer consists of 587 beads in the WT cell and 586 beads in the DELC cell, each bead representing a 10 kb genomic region. We started our simulation with a self-avoiding random walk and run in the NVT ensemble for 10^7^ timesteps to attain equilibrium. Contact probabilities from cHiC were used to determine the constraint between connecting pairs. Constraint parameters were selected to get the best correlation between experimental and simulated cHiC contact maps (Figure S1). (See SI, section 1).

We then analyzed the ensembles of 3D configurations of the considered genomic loci to quantify the E-P interactions and extract the parameters for kinetic models (see SI, section 4). A markov chain based kinetic model was parameterized to reproduce the chromatin dynamics, specifically E-P interactions. All enhancers within the selected loci were identified, and the dwell time distributions were computed for each E-P pair using polymer simulation trajectories (Figure S3). Binding and unbinding rates were computed by fitting exponential curves to the distribution data [19] assuming E-P binding and unbinding as a Markovian process (see SI, section 4). The obtained rates were supplied to a kinetic model that predicts the probabilities of enhancers being attached to the sox9 and kcnj2 promoters and the average number of attached enhancers and TFs in a steady state. The expression level of a gene was assumed to be proportional to the average condensate size of its promoter [20]. (See SI, section 5-7).

### Dataset

To validate our model, a 6 Mb genomic region encompassing the sox9-kcnj2 locus of chromosome 11 in E12.5 mouse limb bud cells was chosen to predict gene expression. We have chosen these specific genomic loci as both structural and expression data is publicly available. Experimental cHiC data in WT cells and DELC cells along with transcription changes resulting from the deletion of the CTCF binding domain at the boundary region of TADs were obtained from GEO accession numbers GSE78109 and GSE125294 for WT and DELC cells, respectively [21]. Enhancers were identified by using VISTA [22] and enhancer ATLAS 2.0 [23] databases. We identified 44 enhancers for sox9, which are consistent with the enhancers reported by Despang *et al.* for sox9 in the considered genomic loci. Six additional enhancers (mm628, mm629, mm630, mm631, mm632, and mm2181) were identified for the kcnj2 gene from these databases for the kcnj2 gene.

## 3. Results

### 3.1. Polymer modeling reveals spatial conformation of chromatin and physical characteristic of TAD arrangement

To generate the three-dimensional (3D) conformation of a given segment of chromatin, we generate a series of polymer confirmations from the MD simulation trajectory (Figure 1a). We repeat the MD simulations with 200 different initial conditions to capture different possible conformations and minimize initial condition bias. In total, 4×10^7^ distinct configurations were recorded to represent the ensemble of 3D configurations of the chromatin segment under investigation. To assess the quality of the simulated 3D structure with the actual chromatin organization within the nucleus, we calculated the Pearson correlation coefficient between the model-derived contact map and the experimental cHiC contact map. The Pearson correlation coefficient of 0.96 indicates a strong correlation (Figure 1c) between the simulated 3D structure and the experimentally measured chromatin structure. Such polymer models have demonstrated their efficacy in elucidating the spatial conformation of chromatin from HiC contact maps [12], [14], [15], [24]–[26].

**Figure 1:**
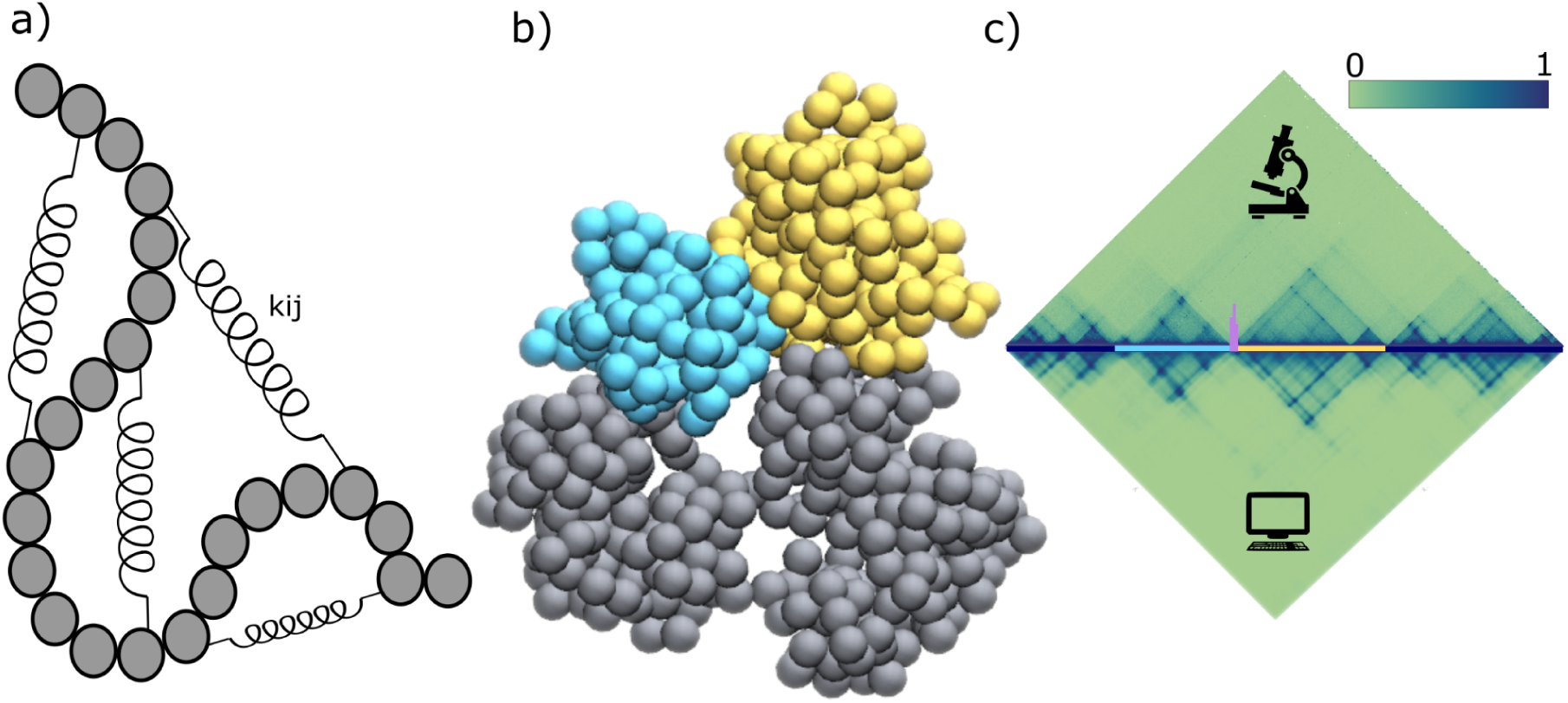
**a)** Schematic representation of polymer model of chromatin is represented by chain of beads where each bead represents 10 kb genomic region and the spring represents the spring constant between two beads; **b)** Time snapshot of the 3D structure of WT cell (cyan and yellow globular domains show the kcnj2 and sox9 TADs, respectively, and silver beads show other genomic regions of the considered 6Mb region (drawn using VMD [30]); **c)** The contact matrices derived from cHi-C experiments performed in E12.5 limb buds (top) and the polymer model (bottom) for WT cell (Pearson correlation = 0.969). Schematic of the genomic region highlights kcnj2 TAD (cyan line) and sox9 TAD (yellow line) separated by CTCF boundary (purple marker).

The generated ensemble of 3D chromatin structure was systematically analyzed to identify distinctive characteristics of the three-dimensional organization of the genomic loci. Our observations revealed the formation of structures resembling globules for each kcnj2 and sox9 TAD. The beads corresponding to a TAD form a single globule and form dynamical connections with other beads in the globule (Figure 1b). Furthermore, beads representing the kcnj2 and sox9 TADs exhibited a striking non-mixing behavior relative to the mixing of intra-TAD beads, such that TADs maintain their respective independent spaces within the 3D conformation.

To correlate gene regulation and chromatin structure, spatial interactions of encoded regions with regulatory elements were investigated. In particular, an exhaustive list of all enhancers encoded in this chromatin segment and associated with sox9 and kcnj2 was created [22], [23]. Subsequently, a focused analysis of enhancers and promoter interactions from the simulated ensemble of configuration was carried out. Our investigations revealed that all 44 enhancers associated with sox9 as identified experimentally by Despang *et al.* were situated within the confines of sox9 TAD. In contrast, the kcnj2 locus resided in an adjacent TAD, spatially separated from all the enhancers. This elucidates the role of TAD boundaries as insulating elements that restrict the interactions between enhancers and promoters located in neighboring TADs [27]–[29].

### 3.2. MD simulation trajectories quantify three-dimensional E-P interactions and highlights the role key-enhancer in gene transcription

Various experimental studies using techniques including 3C, 4C, 5C, and ChIA-PET, have demonstrated that all active enhancers form physical contacts with promoters via chromatin looping or some form of tracking mechanism, despite genomic separation from the promoter [31]. Given that our modeling approach captures the statistical fluctuations of chromatin structure around the HiC data through temporal evaluation of configuration, we can quantify interactions of enhancers and promoters in 3D space and infer the kinetics of these contacts.

To quantitatively investigate the nature of E-P contacts inside the nucleus, we selectively look into the pairwise contacts between 44 enhancers and promoters corresponding to both genes namely, sox9 and kcnj2. These enhancers were located at distances ranging from tens to hundreds of base pairs from the promoters. Our analysis reveals that enhancer beads form a dynamic cluster around the promoter bead, with enhancer beads continuously moving in and out of contact range (1.5σ) of the promoter bead. We track the total number of enhancers surrounding each promoter at a given time and its variation with time. The average number of enhancers surrounding the sox9 is 3.21, with a median value of three (Figure 2a). On the other hand, the kcnj2 promoter in the neighboring TAD is in contact with only one enhancer with a miniscule contact probability of 0.01 (Figure 2a). A closer look reveals that among 44 enhancers, only four enhancers namely E1 (chr11 : 111519300 - 111520200), E34 (chr11 : 112671913 - 112672982), E40 (chr11 : 112982498 - 112983853) and E41 (chr11 : 113008929 - 113009857) have significant contacts with the sox9 promoter, and other enhancers form relatively intermittent contacts with the promoter (Figure 2b). However, the number of enhancers coming in close contact with the sox9 promoter fluctuates and occasionally can reach up to 14 (Figure 2a). The probability of a specific number of enhancers being in contact with promoters is shown in figure 2a.

**Figure 2:**
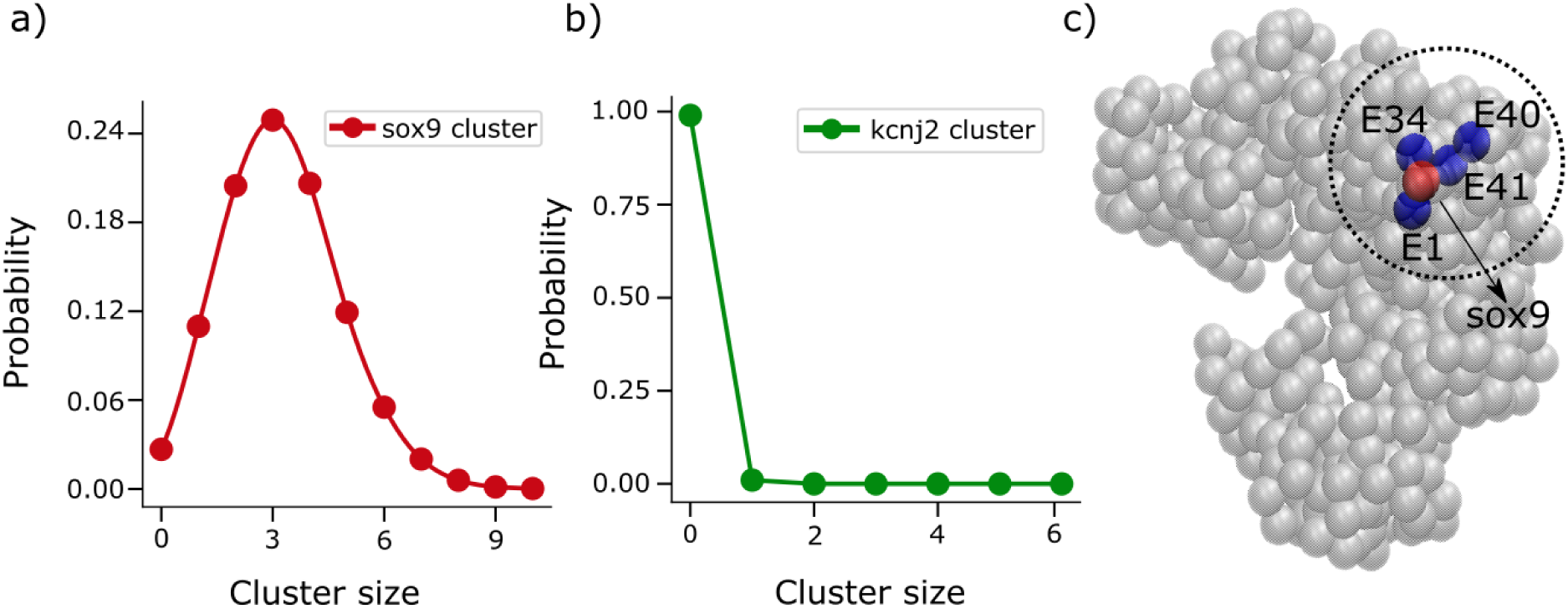
The probability distribution for different sizes of enhancer cluster computed from MD trajectories around **a)** sox9 and **b)** the kcnj2 promoter in the WT cell; **c)** The polymer snapshot (drawn using VMD [30]) shows the enhancers E1, E34, E40, and E41 (blue beads) around the sox9 promoter (red bead).

To identify the most effective enhancers out of the 44 enhancers associated with sox9, we conducted an analysis of the binding affinities between individual enhancers and the sox9 promoter. Among the 44 enhancers examined, E41 has the highest affinity for the sox9 promoter as it spends 26.87% of time in contact with the sox9 promoter, out of which it spends 2.9% of time as a single with the promoter. It spends 18.7% of its time as a diad and 50.1% of its time as a triad (Table I). Similarly, E1, E40, and E34 exhibit 24.98%, 18.55%, and 16.5% time in contact with promoters, respectively. E1 spends 4.4% of its time as a single, 20.8% of its time as a diad, and 42% of its time as a triad out of all the time it attaches to the promoter. Moreover, our analysis also suggests that E34 and E40 spend 4.1% and 4.38% time as singles, 22.5% and 22.2% time as diads and 34.3% and 49.8% time as triads, respectively, out of the time they attach themselves to the sox9 promoter. Other enhancers have a smaller chance to be in contact with the sox9 promoter (0.15%-12.74%). Based on this analysis, we propose that these four enhancers are most important to effectively regulating the sox9 gene expression. The findings of this study indicate that the effective enhancers exhibit cooperative regulation with other enhancers for gene regulation.

**Table I:**
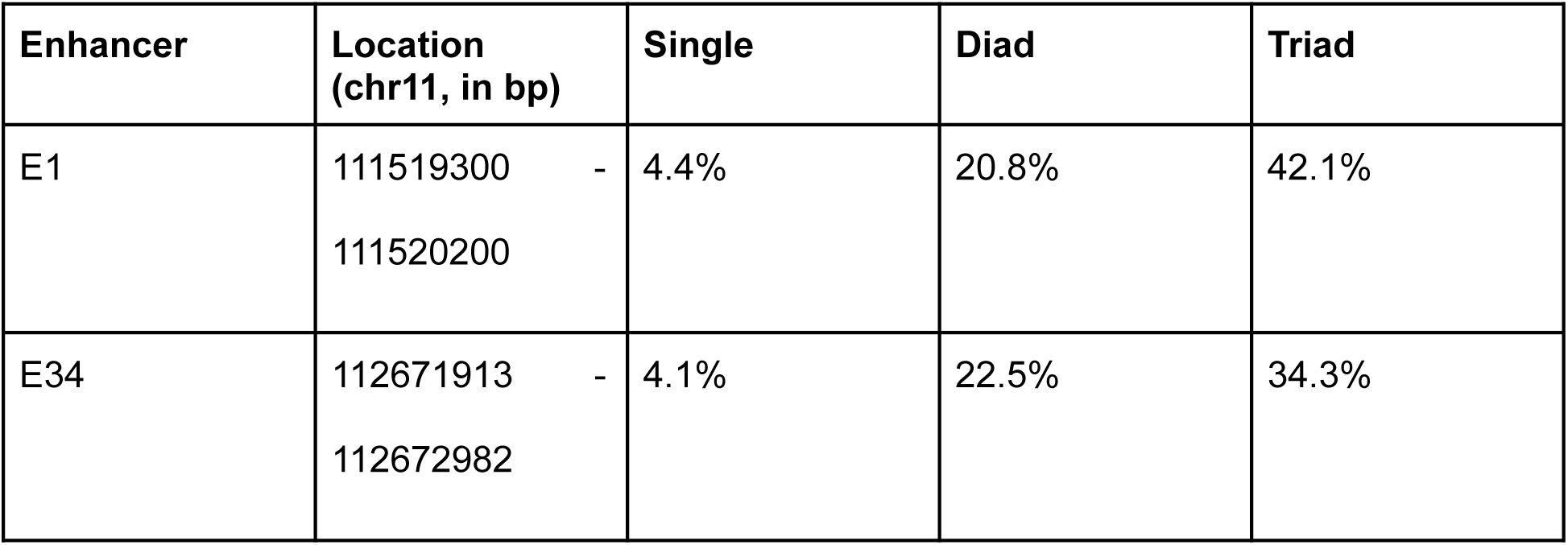
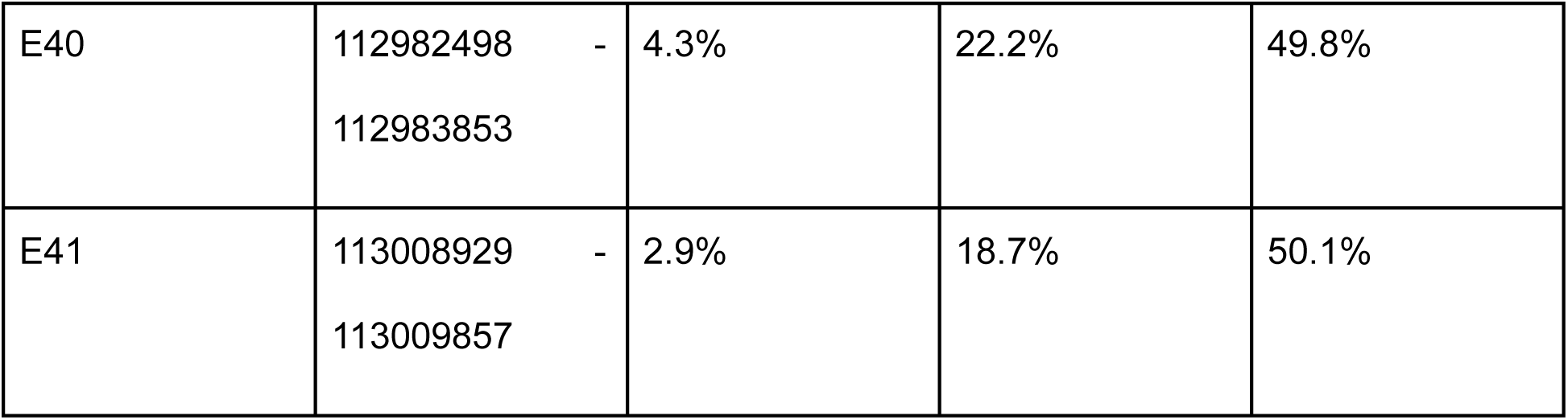
Cooperative nature of enhancers. Probability of the enhancers to be in simultaneous contact with other enhancers.

### 3.3. CTCF TAD boundary deletion leads to TAD fusion, resulting in significant alterations to enhancer-promoter interactions and the spatial organization of enhancer elements

Disruption of chromatin organization can have significant consequences on gene regulation, as it can result in the occurrence of ectopic interactions between enhancers and their target promoters [29]. Despang *et al.* have shown that deletion of all four CTCF binding domains present at the boundary of sox9 and kcnj2 TADs in chromosome 11 of E12.5 mouse limb bud cells leads to the merging of both TADs. Expression analysis reveals that the deletion of major CTCF binding domains leads to a two-fold increase in the kcnj2 expression level, whereas the sox9 expression level decreased by 20% compared to WT cell. Our goal is to reproduce this expression change through our computational framework and then understand how the modulations in E-P interactions give rise to such changes in gene expression. The analysis of molecular dynamics trajectories of chromatin provides a valuable tool for investigating alterations in the dynamics of regulatory elements arising from such structural disruptions. To this end, we simulate the CTCF boundary deletion region between sox9-kcnj2 TADs. Using the same methodology employed for the WT cell, we constructed an ensemble of 3D conformation of the DELC cell (Figure 3a). The contact matrix generated from this ensemble of configurations is highly correlated with the cHiC contact map with Pearson correlation coefficient of R = 0.92 (Figure 3b). The examination of the 3D conformation reveals that in the DELC cell, the two distinct globules representing individual TADs encompassing the sox9 and kcnj2 genes in the WT cell, are merged into a single globule (Figure 3a).

**Figure 3:**
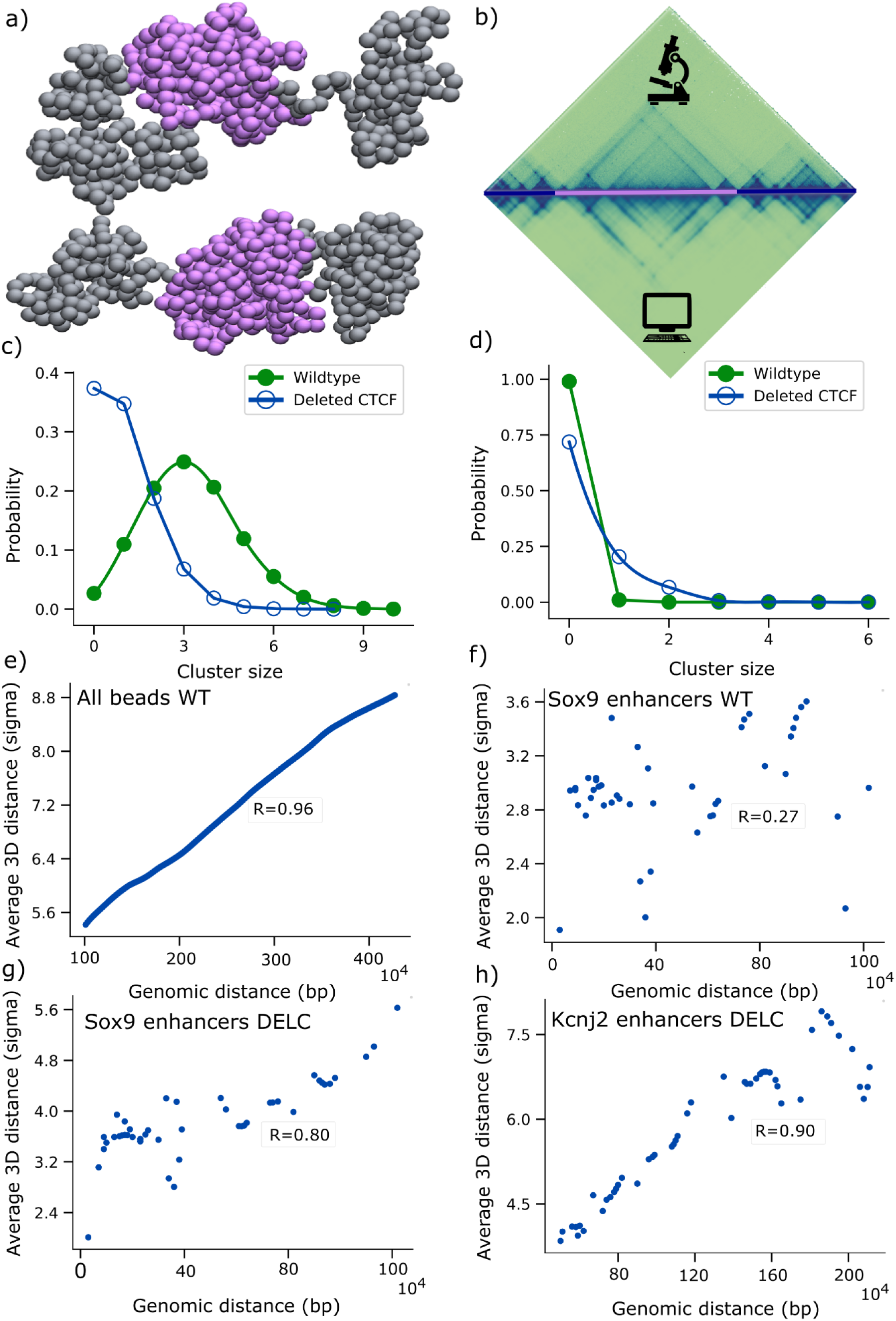
**a)** A representative 3D configuration of DELC chromatin segment cell (Top and bottom structures represents front and back of the 3D ensemble) (drawn using VMD [30]); **b)** The contact matrices derived from cHiC experiments performed in E11.5 limb buds (top) and the polymer model (bottom) for DELC cell (Pearson correlation = 0.92). Purple line in the genomic region highlights merged TAD shown in a); **c)** and **d)** The probability distribution curve of enhancer clusters of different sizes around the promoters shows the enhancer clustered around the sox9 and kcnj2 promoters, respectively. The green curve shows the probability distribution plot for the WT and the blue curve for the DELC cell; The scatter plot between genomic distance (in base pairs or bps) and average 3D distance from the promoters (in sigma units) for **e)** all beads, **f)** enhancers of sox9 gene in WT cell, **g)** enhancers of sox9 gene in DELC cell, and **h)** enhancers of kcnj2 gene in DELC cells, respectively. Here, R represents the Pearson correlation coefficient.

The merging of TADs also impacts the E-P interaction. Based on the ensemble of 3D structures generated using the MD simulation from cHiC data, we compared the E-P interaction for both the cases (WT and DELC). To quantify the changes in E-P interactions resulting from the TAD fusion, we compared enhancer cluster size surrounding both the genes in WT and DELC cells. The average number of enhancers forming clusters around the sox9 was 3.21 before deletion while after deletion this number reduces to 1.04, signifying the reduced accessibility of enhancers post CTCF boundary deletion. On the other hand, the average number of enhancers for kcnj2 increases from 0.001 to 0.36 indicating increased contact with enhancers upon TAD fusion.

To get more insight into the E-P interaction, we compared the distribution of the size of the enhancer cluster before and after CTCF boundary deletion for both the genes. Enhancer cluster size distribution of sox9 shows that E-P interaction significantly decreased for the sox9 gene after TAD boundary deletion (Figure 3d), as the peak of the probability distribution curve corresponding to the size of enhancer-cluster around sox9 promoter shifts from 3 to 0 (Figure 3c). On the other hand, the probability peak remains at 0, but the magnitude of the probability increases from 0.01 to 0.203 in the DELC cell (Figure 3d). Next, we ask the question: are genomic distance from promoters and average physical distance with promoters correlated? To answer this, we calculate the average spatial distance for different genomic distances. We observed a significant correlation (R = 0.96) between the genomic distance and average physical distances between different chromatin regions and sox9 promoters within a TAD (Figure 3e). This correlation is similar to normal polymer behavior, which follows a scaling law between monomer separation and spatial separation [28], [32]. This behavior imposes constraints on genomic regions, effectively preventing the interaction between the distal loci within a TAD. However, interestingly, our analysis reveals that enhancers do not show any correlation between their genomic distance and physical 3D distance from their promoters (Figure 3f). These results highlight that the positions of enhancers within the genome are highly adaptable and context-sensitive in their role of controlling gene expression, ensuring that genes can be activated even if they are far apart on the DNA strand.

Contrary to our findings in WT TADs, in DELC, we observed a strong positive correlation (R _(Sox9)_ = 0.80, R_(Kcnj2)_ = 0.90) between the genomic and physical distances of the enhancer and the promoter for both the genes, sox9 and kcnj2 (Figure 3g and 3h). This implies that after TAD boundary deletion, enhancers lose specific arrangement inside the TAD, and exhibit the relationship similar to the other non-specific genomic regions. This revelation highlights the profound impact of TAD boundaries on orchestrating E-P interactions.

In addition to these observations, we find that enhancers are affected differently depending upon genomic distances from the TAD boundary, despite their co-localization within the same TAD. Notably, the average spatial distances between the enhancers and the sox9 promoter, located near the TAD boundary (e.g., mm627 at chr11 : 112671913 - 112672982, E1 at chr11 : 111519300 - 111520200, E2 chr11 : 111546750 - 111548800, hs1467 chr11 : 111689219 - 111690538), were found to be approximately doubled in the DELC cell compared to the WT cell. On the other hand, enhancers located far away from the boundary exhibit relatively small (1-26%) changes to the average spatial distance with sox9 promoter. This intriguing finding implies that the 3D chromatin conformation, in conjunction with genomic position, plays a pivotal role in governing E-P interactions and, consequently, in gene regulation.

### 3.4. Parameterization and validation of E-P Kinetics

The observed changes in E-P contacts are likely to be responsible for the transcription changes observed in the experiment. However, quantitative correlation and predicting changes in transcription levels due to E-P interaction modifications remain challenging. As gene expression is reliant upon the E-P interactions, we quantify the binding and unbinding of different enhancers with a given promoter from the MD simulation trajectories. In particular, we calculated the binding and unbinding rates of all identified enhancers with both the promoters, as these rates influence the transcription level of the target promoter [33]. The rates were calculated by fitting an exponential function to the dwell time distribution in each of the states of a given E-P pair which were computed from MD simulation trajectories (Figure 4a,b). (See SI, section 4 and Figure S3).

**Figure 4:**
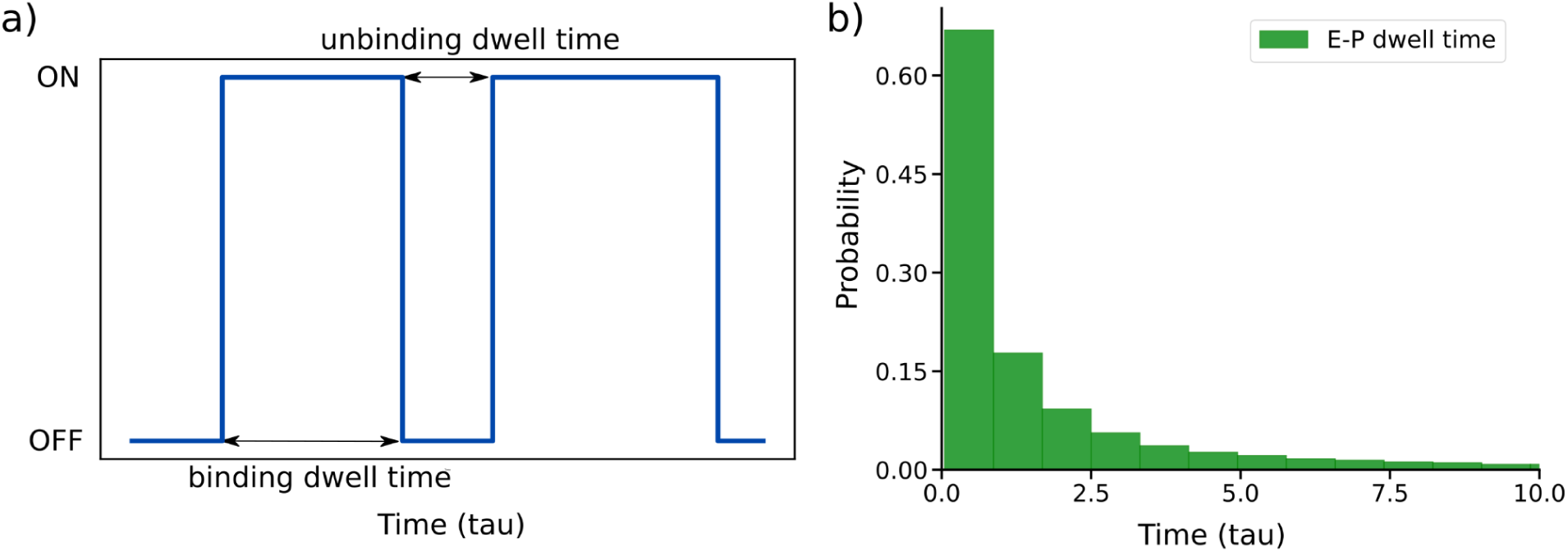
Dwell time distribution of enhancer (E1) bead with sox9 on promoter bead pair: **a)** The schematic shows binding and unbinding of one enhancer with its target promoter. The time is in tau (reduced) units; **b)** The histogram plot of sox9 promoter and E1 enhancer shows an exponential distribution.

We observed that binding/unbinding rates exhibit dependence on the presence of enhancer already paired with promoter. To quantify this, we compute the binding and unbinding rates for a given enhancer for different E-P cluster sizes. We observed a linear decrease (increase) in the binding (unbinding) rates with increasing cluster size (Figure S4). This indicates that it becomes progressively difficult for enhancers to bind a promoter site as E-P cluster size increases. Thus, the number of enhancers bound to a promoter cannot go beyond a threshold number, determined by the thermodynamics of E-P binding. To further understand the dynamics of E-P interactions responsible for the expression of sox9 and kcnj2 genes, we developed a stochastic kinetic model of gene expression. The model specifically accounts for the binding and unbinding of each enhancer with the promoters of both genes (see figure 5a for details). We simulate the model using Gillespie’s algorithm. The input rate parameters for the simulations are computed using MD simulation trajectories (see the Methods and SI, section 4 for details). Using our kinetic model, first we validate if it (Figure 5a) faithfully captures the E-P interaction simulated through our constrained polymer model. To this end, we compute the average E-P cluster size from 100 independent simulation trajectories produced by our kinetic model and compare the distribution of E-P cluster size from both simulations. As shown in the figure 5b, cluster size distribution of the WT sox9 gene matches quite well with the polymer simulation model. This shows that our kinetic model can accurately capture the E-P dynamics.

**Figure 5:**
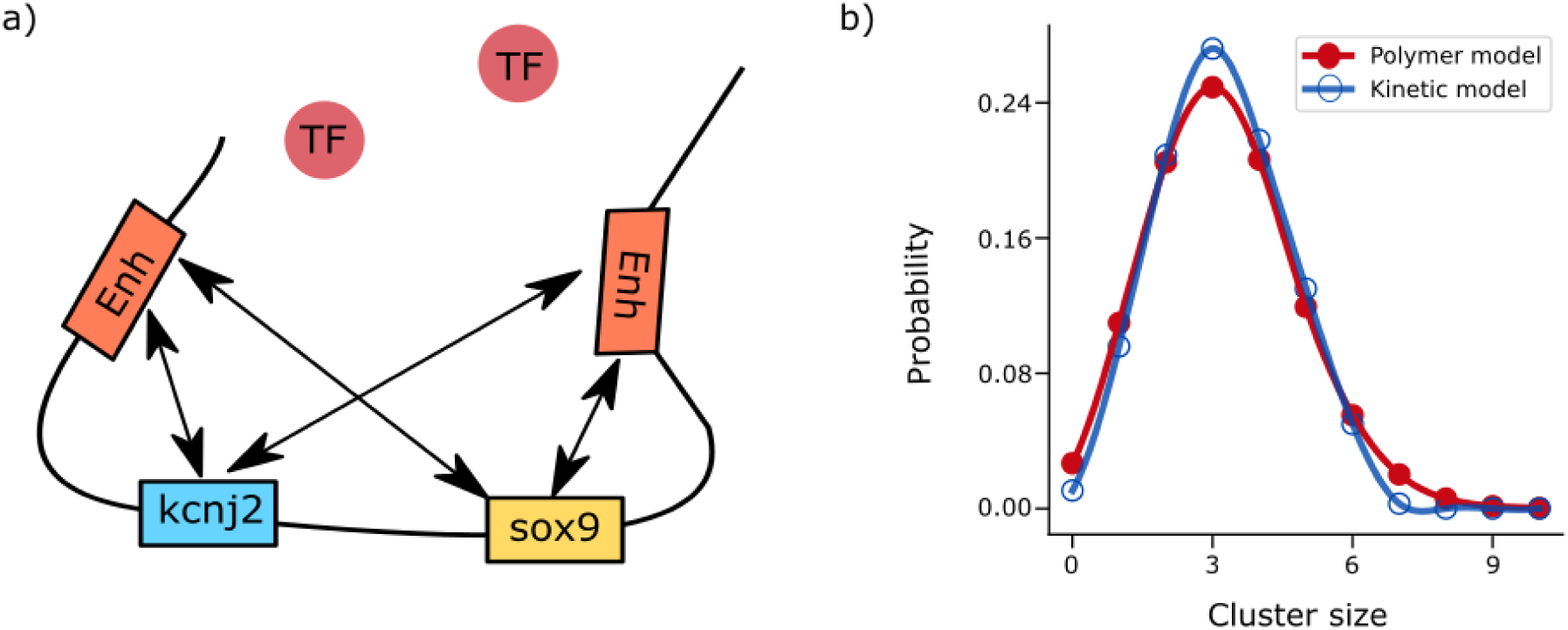
**a)** Schematic of the kinetic model: kcnj2 and sox9 promoters can bind to enhancers with their respective binding and unbinding rates on the genomic location. The transcription factors are represented by spheres which move freely in the nucleoplasm and can bind /unbind with both the promoters; **b)** validation of the kinetic model. The probability distribution plot of enhancer cluster size around the sox9 promoter in polymer simulation (red curve) and kinetic model (blue curve) is shown.

### 3.5. Markov chain integrating E-P interaction and TF binding predicts the transcription changes in the structural variants

While E-P binding plays a crucial role in transcription regulation, TFs also participate in the process. Xiao and co-workers developed a kinetic model based on the E-P and TF-promoter binding rates to predict the transcription level of a gene. We employ a similar approach to predict the level of transcription for a given set of E-P binding/unbinding rates. In this analysis, E-P binding and unbinding rates were computed using the simulation trajectories, whereas TF binding and unbinding rates were chosen to be the same as provided in the kinetic model of Xiao and coworkers [20].

We incorporate the influence of TF as an independent binding/unbinding event (see SI, section 5). In the model, we assumed that the expression level of a gene is directly proportional to the average number of enhancers and weighted number of TFs (cluster) attached to the promoters [34]. We use the same weight factor for both WT and DELC cells. Remarkably, we found that the relative decrease in gene expression levels of sox9 measured in the WT cell compared to the DELC cell matches the findings of the experiments by Despang *et al.* aforementioned study (Figure 6). Our model was able to quantitatively predict the gene expression changes due to TAD boundary deletion. We could directly connect the structural changes with the gene regulation through this approach. Our approach not only gives a simple tool for predicting gene expression but also provides insights into the impact of E-P interaction on gene regulation.

**Figure 6:**
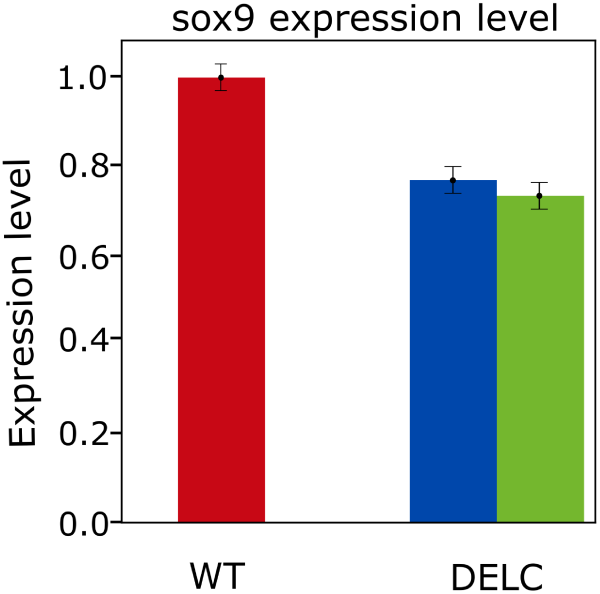
Relative expression of sox9 in E13.5 limb buds for WT and DELC cells. Bars represent the mean of condensate size and error bars represent the standard deviation. The WT gene expression is normalized to one both for experiment and simulation. For the DELC cell, the blue bar shows the experimental result and the green bar shows the simulation result.

### 3.6. Newfound enhancer-accessibility drives large change in gene expression level of kcnj2 gene post boundary deletion

To understand the remarkable increase in the kcnj2 gene expression levels in the DELC cell with respect to the WT cell, we closely examine contacts of the kcnj2 promoter with various enhancers reported in different studies [22]. We identified a total of nine possible enhancers specific to kcnj2. Out of these nine, six enhancers (mm628, mm629, mm630, mm631, mm632, and mm2181) affect only kcnj2, and the three enhancers (mm627, mm634, and mm1285) shared by both sox9 and kcnj2. We refer to these nine enhancers as kcnj2-specific enhancers. Through MD trajectory, we observed that all kcnj2-specific enhancers, which were identified from the Vista enhancer browser, were present in the sox9 TAD in the WT cell [22] (Figure 7a) and rarely made contact with the kcnj2 promoter. However, they form a cluster around kcnj2 and frequently come in contact with the promoter after TAD boundary deletion (Figure 7b). This suggests that these enhancers may not be active in WT of this particular cell type and might have been encoded to be useful in alternate chromatin configuration corresponding to other cells. Our findings also suggest that structural elements such as the TAD boundary can be instrumental in enhancers selectivity [35].

**Figure 7:**
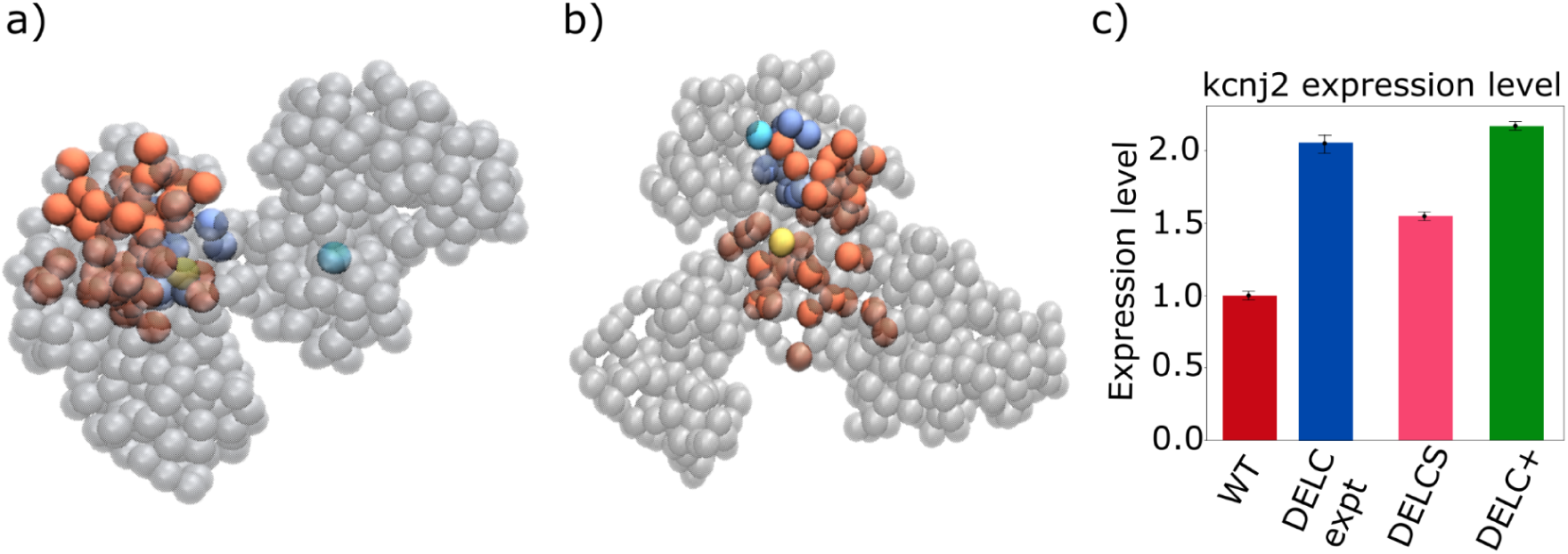
3D configuration of chromatin segment captured from MD trajectories highlighting the representative positions of enhancers with sox9 and kcnj2 promoters in **a)** WT cell and **b)** DELC cell. Sox9 promoter, kcnj2 promoter, specific and non-specific enhancers are represented by yellow, cyan, blue and orange beads, respectively, and other beads are made transparent (drawn using VMD [30]); **c)** Relative expression of kcnj2 in E13.5 limb buds for WT and DELC cells. DELCS and DELC+ bars show the relative expression of kcnj2 considering only the specific enhancers and considering both specific and non-specific enhancers, respectively, and error bars represent the standard deviation over 100 initial conditions. The WT gene expression is normalized to one both for experiment and simulation.

Next, we examine if the E-P interaction with the kcnj2-specific enhancers is responsible for enhanced transcription of kcnj2 in DELC. In other words, the evidence of E-P contact in spatial arrangement of chromatin can be attributed to the functionality of an enhancer. We propose to use the spatial contact information about E-P interactions easily obtained from the simulated 3D genome structure, as a computational tool for identifying active enhancers in a given chromatin conformation. To quantify the impact of these specific enhancers, we computed the relative contributions of the specific and nonspecific enhancers in the expression (Figure 7c). Our results show that specific enhancers contribute to nearly 70% of the total gene expression in the DELC cell (Figure 7c), thereby, the significant increase in the kcnj2 expression can be attributed to pairing with specific enhancers. Based on this, we believe that physical distances between enhancer and promoters, extracted from MD simulations, can be used to identify the active enhancers.

## 4. Discussions and Conclusions

Our study introduces a computationally efficient approach to predict gene expression based on the spatial organization of chromosomes derived from cHiC contact maps. By conducting a systematic analysis of 3D chromatin organization, we unveil specific folding patterns within the chromosome conformation (Figure 1c and 3b). Our polymer model was able to show the changes of E-P interaction due to genome rearrangements like TAD boundary deletion (Figures 3c and 3d). Our model also gives insights into the role of chromatin conformations into determining the specificity of E-P interaction (Figure 7). The temporal evolution of E-P contacts extracted from trajectory was shown to be a good indicator of the functionality of an enhancer. Several modeling studies have yielded valuable insights into the fundamental aspects of chromatin organization. However, these studies do not make quantitative predictions of gene expression for different chromatin conformations and gene regulation [12], [14], [24], [25]. In contrast, our framework offers a highly effective approach to quantify relative gene expression for different chromatin conformations. This quantification helps to develop an understanding of the intricate relationship between chromatin conformation and the regulation of gene expression. By taking a specific case of structural disruption through TAD boundary deletion, and predicting the transcription changes consistent with experimental results, we validate the ability of our model to make quantitative prediction of gene expressions. Our study introduces a simple framework that captures the key features of structure-dependent gene regulation, such as alteration of E-P interactions resulting from significant CTCF deletions across TAD boundaries, and accessibility of the promoters for TF binding. The present analysis underscores the significance of the spatial genomic architecture in governing the interaction between enhancers and promoters [36]–[38].

Our finding also uncovers the impact of specific chromatin folding on establishing the enhancer specificity. Specifically, we demonstrate that a few enhancers specific to kcnj2 are inaccessible to promoter sites due to presence of the TAD boundary, and are able to make contacts after the deletion of TAD boundaries. Among the nine enhancers identified computationally, only three enhancers, namely mm627, mm628 and mm634, have been experimentally validated as kcnj2 enhancers in these cells [22]. Our analysis confirms that only these enhancers are capable of forming contacts with kcnj2 promoter, but are restricted by the specific genome folding achieved with the help of TAD boundaries. However, deletion of the TAD boundary removes these topological constraints, and E-P contacts are possible (Figures 7a and 7b).

Although we have used our model to predict the change in transcription due to deletion of the TAD boundary in limb bud cells, it can be easily extended to other cell types and means of chromatin structural changes. The applicability of our model is limited only to the availability of HiC data. As long as HiC data is available, our model can be used to compare the relative changes in transcription in different chromatin organizations. We envision that our model will be particularly helpful in predicting the changes in transcription due to nuclear deformation arising under various biological scenarios and in understanding the role of chromatin organizational changes during the cell development cycle and cell fate decisions.

We briefly summarize the underlying assumptions of our model, although crucial to the model’s architecture. Firstly, our model considers chromatin dynamics within nucleoplasm which exhibits viscosity ranging from 0.03 P to 0.1 P [12], [13], [25] which may depend on several factors such as the state of cell and cell type. To understand how viscosity may affect our final results, we simulated the specified genomic region for a range of viscosity values and were able to reliably reproduce E-P cluster size distribution for all viscosity values (Figure S2). It should be noted that viscosity affects the relaxation dynamics of the chromatin and short-term behavior may still depend on the viscosity values. Furthermore, each bead corresponds to a 10 kbp genomic region in our model, aligning with the resolution of experimental cHiC contact maps. However, it is important to acknowledge a resolution discrepancy between our model and the physical distribution of enhancers and promoters, which often span a smaller genomic locus with an average value of 50-1500 bps [1]. Despite this disparity, we can overlook this limitation, as for this particular genomic region, each enhancer falls in separate beads of the polymer model. An avenue for model improvement lies in increasing the polymer model’s resolution to better align with these finer genomic details, which again depend upon the resolution of the Hi-C experiment. In addition to that, in our kinetic model, we assumed uniform transcription factor binding rates for both promoters and enhancers due to the absence of detailed data and for the sake of model simplifications. While TF’s binding may indeed vary across different enhancers and promoters, it is also true that there are often common motifs or binding sequences that certain transcription factors recognize. In such cases, assuming equal binding and unbinding rates is a reasonable approximation when modeling the system at a coarse-grained level.

In summary, we have developed a computational method to predict the gene expression corresponding to the cHiC map. Through the utilization of this modeling technique, the complex regulatory network’s dynamic interactions are effectively captured. This not only contributes to the advancement of our fundamental comprehension of genome architecture but also offers significant insights into potential therapeutic interventions that aim to address abnormal gene regulation in pathological circumstances.

## Supporting information

Supplementary details of methods

## Acknowledgement

HK acknowledges the financial support from the Department of Science and Technology, India under the “Fund for Improvement of S&T Infrastructure (SR/FST/PS-I/2020/140)” scheme.

## Notes

### Competing Interest Statement

The authors have declared no competing interest.

