## Supplementary details of methods for "Polymer Model Unveils Quantitative Association of Chromatin Conformation and Gene Regulation"

### 1. Molecular dynamics simulation:

To model the loci of interest, we represented each bead in the bead-spring model as a 10 kb genomic region. The polymer model consists of 587 beads in the wildtype cell and each bead is connected to the next bead with FENE potential with with a distance cutoff of 1.5σ and strength of 30 $k_{\beta}T/\sigma^2$. The non-overlapping of beads is implemented through the lennard-jones interaction interaction between all beads with an interaction cutoff distance of 1.12σ. To enforce chromatin conformation on the polymer model, we apply harmonic constraints between different beads such that magnitude of the spring force constant is proportional to contact probabilities from HiC map. Different power law scalings were tested to determine the optimum relation between contact probabilities and force constant k. Our calculations suggest that $k \propto c^2_{ij}$ generates the conformations which exhibit the best correlation with HiC (Figure S1).

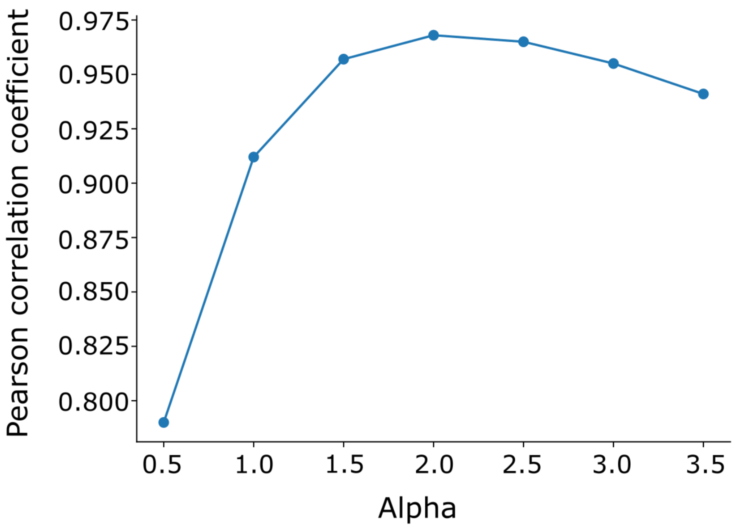


***Figure S1:*** *Parameterization of $C_{ij}^{\alpha}$. The X-axis shows the variation of $\alpha$ and the Y-axis shows corresponding Pearson correlation coefficient.*

We used the LAMMPS package to model the bead spring polymer [1]. The equation of motion is integrated using the Verlet algorithm and temperature is kept constant using Langevin thermostat (T=1.0) [2]. The nucleoplasm was modeled as a viscous fluid with friction coefficient of a particle $\zeta$ = 1.0 in reduced units surrounding the polymer. To ensure that our key results are not affected by the choice of parameters, we have reproduced key results with five different values of nucleoplasm viscosity (Figure S2) [3], [4].

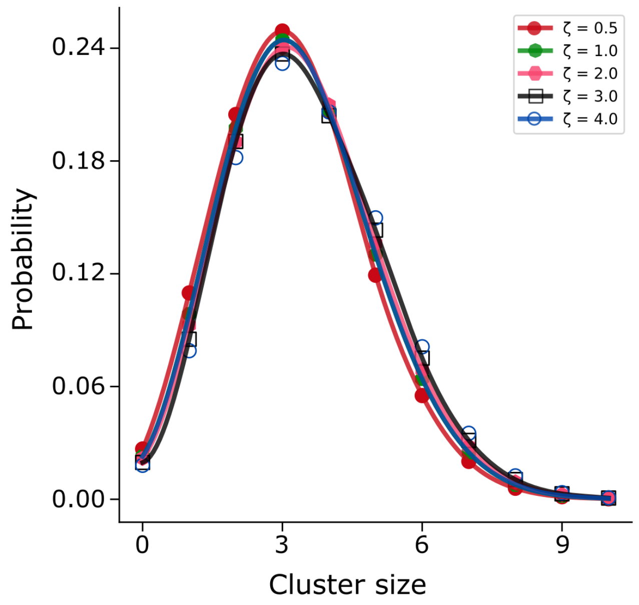


***Figure S2:*** *The probability distribution of enhancer cluster size for different friction coefficient values.*

### 2. Loci and datasets:

To validate our approach, we choose the 6 Mb genomic region of sox9-kcnj2 locus of chromosome 11 of E12.5 mouse limb bud cells. Experimental study has demonstrated the TAD-merging and transcription changes of genes enclosed in this region, namely sox9 and kcnj2 after targeted deletion of CTCF binding domain (chr11: 109010001 – 114880000, mm9). We have modeled both cells and refer to them as wildtype (WT) and CTCF boundary deleted (DELC) cells throughout the manuscript. The cHiC kr-normalized 10-kb resolution data was taken from GEO accession numbers GSE78109 and GSE125294 [5] for WT and DELC, respectively.

### 3. Conversion of reduced units to physical units:

For sake of simplicity, we have performed the polymer simulations in reduced units. We set the diameter of each bead in the polymer simulation to $\sigma$ and take it as unit length, and the mass of each bead to be unit mass. The energy scale unit is considered as $\epsilon = k_\beta T$. The physical diameter of the bead $\sigma$ is estimated by following the same approach as [2] and found it to be 75 nm. The standard MD approach is used to map dimensionless parameters into physical units. For instance, if we take viscosity to be 0.1P and temperature to be 300K, then the time unit can be calculated to be 0.1 sec by using the MD relation $\tau = \eta(6\pi\sigma^3/\varepsilon)$. We used a time integration step of 0.001 in the MD simulation and let the system evolve up to $10^7$ timesteps. We performed 200 individual MD simulations for each set of system parameters.

4. **Analysis from three-dimensional polymer structure:**

In this study, we utilized nearly 200 molecular dynamics (MD) simulations to investigate the enhancer cluster size distribution of each promoter in both WT and DELC cells. We located all enhancers within the considered loci reported by Despang *et al.* and determined the binding and unbinding dwell times for each enhancer-promoter pair. The binding dwell time refers to the duration for which an enhancer remains bound to a promoter, while the unbinding dwell time refers to the duration for which it remains unbound [6]. We plotted the dwell time distribution curve for each enhancer-promoter pair and calculated the binding and unbinding rates by fitting the distribution curve with a double exponential curve: $y = ae^{-bx} + ce^{-dx}$.

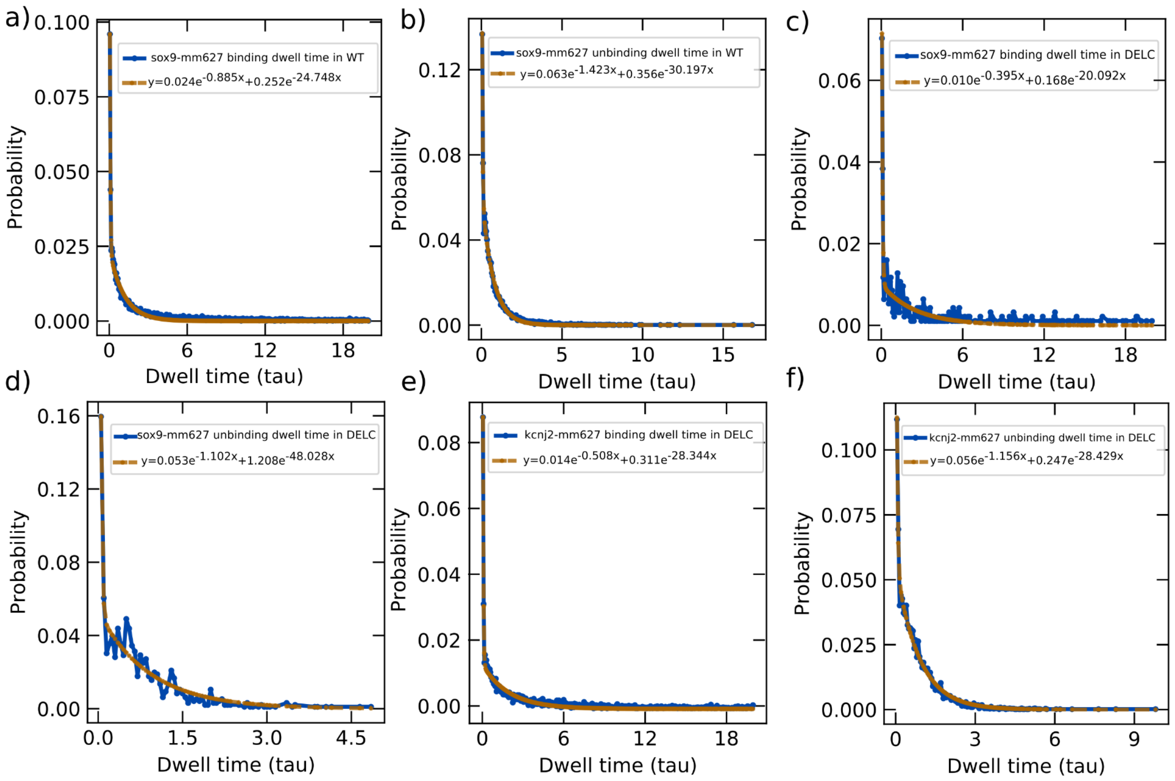


***Figure S3:*** *Exponential fitting of probability distribution of dwell time (in Lennard Jones or LJ units) of enhancer-promoter pairs* ***a)*** *binding of mm627 enhancer with sox9 promoter in WT cell,* ***b)*** *unbinding of m627 enhancer with sox9 promoter in WT cell,* ***c)*** *binding of mm627 enhancer with sox9 promoter in DELC cell,* ***d)*** *unbinding of m627 enhancer with sox9 promoter in DELC cell,* ***e)*** *binding of mm627 enhancer with kcnj2 promoter in DELC cell,* ***f)*** *unbinding of kcnj2 enhancer with sox9 promoter in DELC cell. The solid line represents the simulation dwell time data and the dashed line is the fitted curve.*

We also calculated the cluster-dependent and enhancer-promoter pair independent rates from the cluster size distribution data. To determine the cluster-dependent binding rate, we fitted the exponential curve to the time it takes to transition from one cluster size to the next larger size. Similarly, we fitted the exponential curve to the time distribution of the enhancer transitioning from one cluster size to another in a decreasing order to determine the cluster-dependent unbinding rate. We used linear regression to fit the data obtained and calculate the slope for each cluster size. These analyses were performed for both WT and DELC cells.

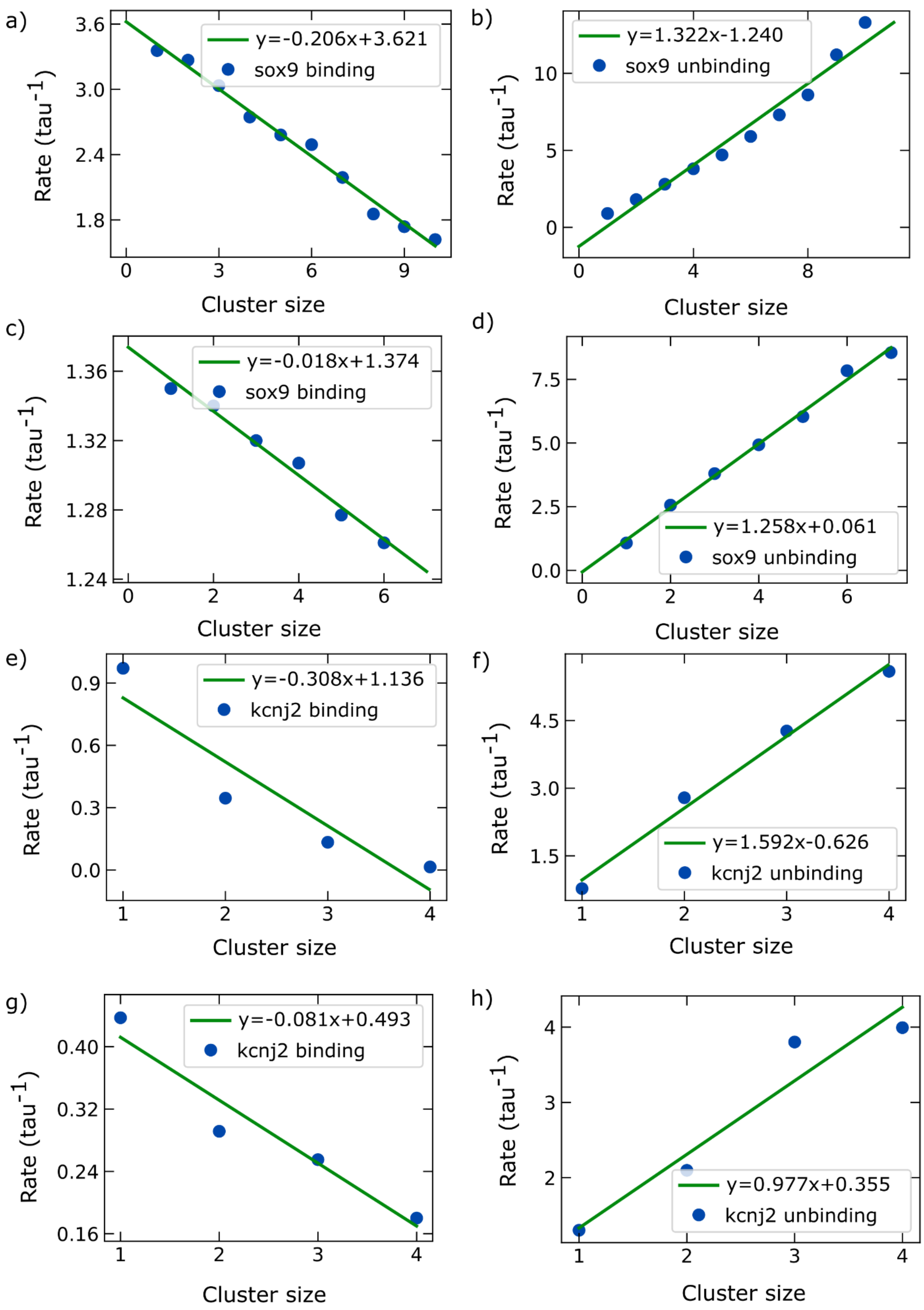

a)
y=-0.206x+3.621
sox9 binding
Rate (tau$^{-1}$)
Cluster size
b)
y=1.322x-1.240
sox9 unbinding
Rate (tau$^{-1}$)
Cluster size
c)
y=-0.018x+1.374
sox9 binding
Rate (tau$^{-1}$)
Cluster size
d)
y=1.258x+0.061
sox9 unbinding
Rate (tau$^{-1}$)
Cluster size
e)
y=-0.308x+1.136
kcnj2 binding
Rate (tau$^{-1}$)
Cluster size
f)
y=1.592x-0.626
kcnj2 unbinding
Rate (tau$^{-1}$)
Cluster size
g)
y=-0.081x+0.493
kcnj2 binding
Rate (tau$^{-1}$)
Cluster size
h)
y=0.977x+0.355
kcnj2 unbinding
Rate (tau$^{-1}$)
Cluster size

***Figure S4: a)** Binding rates of enhancers for different cluster sizes for sox9 in WT cell, **b)** unbinding rates of enhancers for different cluster sizes for sox9 in WT cell, **c)** binding rates of enhancers for different cluster sizes for sox9 in DELC cell, **d)** unbinding rates of enhancers for different cluster sizes for sox9 in DELC cell, **e)** binding rates of enhancers for different cluster sizes for kcnj2 in DELC cell, **f)** unbinding rates of enhancers for different cluster sizes for kcnj2 in DELC cell, **g)** and **h)** binding and unbinding rates of enhancers for different cluster sizes of kcnj2 promoter in DELC cell (considering only specific enhancers), respectively.*

### 5. Stochastic simulation and model parameters

To understand how E-P interaction and the assembly of transcription factors (TFs) at the promoter regulate the gene expression levels in WT and DELC sox9 and kcnj2 genes. We developed a stochastic simulation model (kinetic model) using Gillespie's algorithm. Our model includes two promoters belonging to the sox9 and kcnj2 genes, respectively. The enhancers and TFs can bind to these promoters to start the gene transcription. Enhancers specific to sox9 and kcnj2 genes can bind/unbind to these promoters with rates $e_{ij}/e_{ji}$ These rates are derived from the dwell time distribution of enhancer-promoter interaction in the 3D polymer structure derived from cHiC data (see section 4). Transcription factors bind/unbind to the promoters via direct or indirect mechanisms. In the direct mechanism, TFs in a nucleoplasm diffuse and attach to the promoter of a gene with rate $r$ or it dissociates back into the nucleoplasm with rate $g$. In the indirect mechanism, when the enhancer contacts the promoter, a TF is added to the promoter of a gene and it dissociates into the nucleoplasm through the direct mechanism. In our model, we assume that the direct binding/unbinding rates $r/g$ TFs are independent of the promoter type. The possible number of enhancers and promoters that can bind to the promoter should be finite. Therefore, to prevent an uncontrolled and potentially unrealistic number of enhancers and TFs from attaching to a promoter, we set its limit to $C_{max}$ in our model. The state of an enhancer, is defined as $S_{ij} = \{C_{i,j}: i = [1, 2, ... n] \& j = [0, 1, 2]\}$, where $i$ represents the enhancer number ranging from 1 to n and $C_{i,j}$ is a binary number . The state of an enhancer is classified into three categories: enhancer attached to sox9 ($j = 1$) or kcnj2 promoter ($j = 2$) or is unbound ($j = 0$). Now, the element of the state $S_{ij}$ is defined as, $C_{11} = 1$ implies that the enhancer $e_1$ is attached to the promoter one i.e promoter of the sox9 gene and $C_{22} = 1$ implies the enhancer $e_2$ is attached to the kcnj2 promoter. The time evolution of a probability of an enhancer $i$ being attached to the sox9 and kcnj2 promoter can be defined as

$$\frac{dp_i}{dt} = e_{ji}(n(t))p_j - e_{ij}p_i$$

where $e_{ji}$ depends on the number of enhancers attached to a promoter at time t. To determine the average number of enhancers and TFs attached to a promoter we have used Gillespie's algorithm. The gene transcription is influenced by the E-P interaction as well as the availability of TFs. Assuming a direct proportional relationship between the average number of enhancers and TFs attached to a gene's promoter and its gene expression level, this model enables us to measure the expression level of a gene.

6. **Transcription level determination:**

We assumed that the transcription level was proportional to the condensate size of the promoter [7]. We ran the kinetic model for a long time to find the average condensate size of the sox9 and kcnj2 promoters. Here, we calculate the average size of the cluster by taking the weighted size of the enhancer and TFs, as we assume that enhancers are already bound by TFs and we can use a scaling factor to calculate the condensate size [7], [8].

7. **Specific enhancer detection from databases:**

We looked for possible enhancers through the available databases, VISTA and Enhancer ATLAS 2.0 [9], [10]. We could not find any additional enhancers for the sox9 gene other than the ones reported by Despang *et al* [5]. But for the kcnj2 gene, we found six additional enhancers which are mm628, mm629, mm630, mm631, mm632, and mm2181.
